# The Role of Alcohol Consumption on Acetaminophen Induced Liver Injury: Implications from A Mathematical Model

**DOI:** 10.1101/2020.07.07.191916

**Authors:** Aditi Ghosh, Isaac Berger, Christopher H. Remien, Anuj Mubayi

## Abstract

Acetaminophen (APAP) overdose is one of the predominant causes of drug induced acute liver injury in the U.S and U.K. Clinical studies show that ingestion of alcohol may increase the risk of APAP induced liver injury. Chronic alcoholism may potentiate APAP hepatotoxicity and this increased risk of APAP toxicity is observed when APAP is ingested even shortly after alcohol is cleared from the body. However, clinical reports also suggest that acute alcohol consumption may have a protective effect against hepatotoxicity by inhibiting microsomal acetaminophen oxidation and thereby reducing N-acetyl-p-benzoquinone imine (NAPQI) production. The aim of this study is to model this dual role of alcohol to determine how the timing of alcohol ingestion affects APAP metabolism and resulting liver injury and identify mechanisms of APAP induced liver injury. The mathematical model is developed to capture condition of a patient of single time APAP overdose who may be an acute or chronic alcohol user. The analysis suggests that the risk of APAP-induced hepatotoxicity is increased if APAP is ingested shortly after alcohol is cleared from the body in chronic alcohol users. A protective effect of acute consumption of alcohol is also observed in patients with APAP overdose. For example, simultaneous ingestion of alcohol and APAP overdose or alcohol intake after or before few hours of APAP overdose may result in less APAP-induced hepatotoxicity when compared to a single time APAP overdose. The rate of hepatocyte damage in APAP overdose patients depends on trade-off between induction and inhibition of CYP enzyme.

## 1 Introduction

Acetaminophen (APAP) overdose and toxicity is one of the predominant causes of acute liver failure in the United States claiming to take about 500 lives annually (Rumack and Matthew 1975; Paulose-Ram et al. 2003). APAP overdose here is defined as intake of more than 5gm of APAP in one sitting (Remien, Adler, et al. 2012; Remien, Sussman, and Adler 2014). APAP induced hepatotoxicity in patients is believed to be affected by alcohol intake (ethanol, EToH) with some even reporting death due to this liver injury (Lesser, Vietti, and Clark 1986; McClain et al. 1980). Various studies indicate that alcohol regulates the formation of NAPQI by induction of cytochrome enzymes P450CYP2E1, CYP1A2 and CYP3A4 (Lucas et al. 1995; Raucy et al. 1989; Patten et al. 1993). However, the most significant contribution in modulating NAPQI is known to occur through CYP2E1 induction. Alcohol ingestion thus affects NAPQI formation, depending on the amount of alcohol consumption and the time lag between alcohol and APAP ingestion. Thus timing of the intake of alcohol and APAP ingestion is crucial in determining the risk of APAP hepatotoxicity.

It has been suggested in many clinical studies see (Maddrey 1987; Bray et al. 1991; Zimmerman and Maddrey 1995; Draganov et al. 2000) that patients with chronic alcoholism have increased risk of severe hepatotoxicity from single overdose of APAP. Patients with chronic alcoholism are at higher risk to develop APAP-induced liver injury due to induction of CYP2E1 also known as MEOS, the microsomal alcohol oxidizing system, responsible for production of NAPQI from APAP. An elevated rate of CYP2E1 activity amounts to conversion of a greater proportion of APAP to NAPQI and thereby increases the risk of APAP induced liver failure.

Alcohol (Ethanol) also acts as substrate for CYP2 and inhibits the metabolism of other substrates in the enzyme by binding it to the active site (see Chien, Thummel, and Slattery 1997; Thummel et al. 2000; Lieber 1997). For example, clinical studies suggest that in the case of acute alcohol ingestion CYP2E1 inhibits microsomal APAP oxidation leading to decreased NAPQI production. Hence the protective effect of alcohol ingestion due to inhibition of CYP2E1 is limited to the acute case.

Thus, the simultaneous induction and inhibition effect of alcohol on CYP2E1 may play an important role in determining the extent of liver damage in APAP overdose in conjunction with alcohol ingestion. The relative timing may be critical. Alcohol ingestion can both alleviate and aggravate APAP induced liver toxicity depending on the amount of alcohol(captured as Ethanol) consumption and the timing of last alcohol and APAP ingestion (seeBanda and Quart 1982; Rumack, Peterson, et al. 1981; Rumack 1984; Critchley et al. 1983; Schmidt, Dalhoff, and Poulsen 2002). Remien et.al. have developed a mathematical Model for Acetaminophen induced Liver Damage to describe acute liver injury due to one time APAP overdose (Remien, Adler, et al. 2012). The *Model for Acetaminophen induced Liver Damage* (MALD) uses a patient’s aspartate aminotransferase (AST), alanine aminotransferase (ALT), and international normalised ratio (INR) measurements on admission to estimate overdose amount, time elapsed since overdose, and outcome in terms of hepatotoxicity. Here we have extended this modeling framework to incorporate varying level of alcohol consumption and different mechanisms leading to hepatotoxicity. The effect of alcohol on APAP metabolism and hepatotoxicity are captured by mechanisms such as the induction and inhibition effect of alcohol on the CYP2E1 enzyme.

In the present study, the model uses single time APAP overdose and time varying alcohol ingestion to estimate hepatocyte damage. The model also explores APAP overdose patients that are either post/prior (to overdose) acute or chronic alcohol users. The goal of the paper is to highlight how alcohol ingestion impact the progression of liver damage in APAP overdose patients. In particular, the study aims to understand the role of time and frequency of alcohol consumption post a single time APAP ingestion aggravates or alleviate conditions of APAP induced liver injury.

We organize the paper as follows. Section 2 provides a detailed description of the mathematical model. Equilibrium points and stability analysis are discussed in Section 2.2. We also provide a bifurcation analysis analysis with respect to an important parameter here. Numerical simulation of the mathematical model to understand the the extent of liver injury due to single time APAP overdose along with alcohol ingestion at different times is presented in section 3. This model is also simulated for chronic and acute use of alcohol. We conclude with a discussion in Section 4.

## 2 Methods

### 2.1 Model Development and Description

To model the effect of alcohol(captured here by Ethanol, EToH) on APAP-induced liver injury, the model MALD (see Remien, Sussman, and Adler 2014) is connected to a physical pool of CYP2E1 as in (Chien, Thummel, and Slattery 1997). The original model MALD is updated to include the mechanism of alcohol metabolism. APAP is metabolised via glucoronidation, sulfation due to overdoze of APAP and CYP enzymes initiated by alcohol intake. APAP is glucoronidated by hepatocytes at a rate *k_g_*. Sulfation of APAP here follows a Michaellis-Menten kinetics (Ben-Shachar et al. 2012) and induction/inhibition of CYP2E1 by alcohol follows Hill kinetics. Synthesis of enzymes occurs at rate *k_cyp_* and CYP2E1 is cleared at the rate 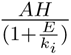 where A is the amount of APAP, H is the number of functional hepatocytes, E is the alcohol amount and *k_i_* is the enzyme dissociation constant. *σ_et_* denotes a constant intake of alcohol over time, it is zero in the case of acute ingestion of alcohol and non zero for chronic ingestion. Alcohol acts as a competitive inhibitor and follows Michaellis-Menten kinetics. The unmetabolised alcohol excreted through urine, sweat or expired air is cleared from the body at a clearance rate *δ*_*E*_. We assume here a single physical pool of CYP2E1 enzyme *C*. As in (Chien, Thummel, and Slattery 1997), CYP2E1 enzyme degenerates rapidly after alcohol is introduced in the body at a degradation rate *k_deg_*. The normalised level of enzyme is denoted by *C*_01_. The time course of the CYP2E1 enzyme *C* is given by equation 6.

*Total Body APAP*:

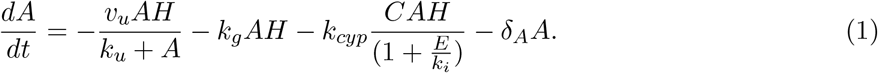

*Intracellular NAPQI Concentration*:

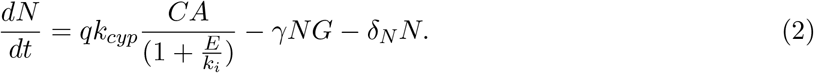

*Intracellular GSH concentration*:

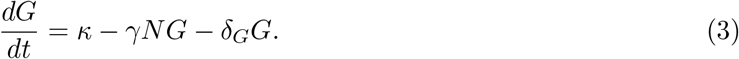

*Functional Hepatocytes*:

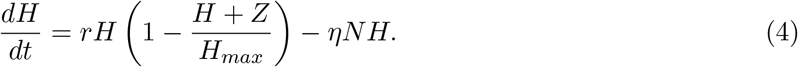

*Damaged Hepatocytes*:

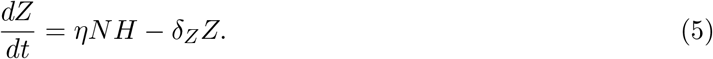

*CYP2 Enzyme Concentration*:

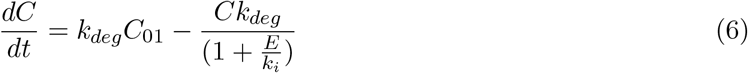

*Alcohol Concentration*:

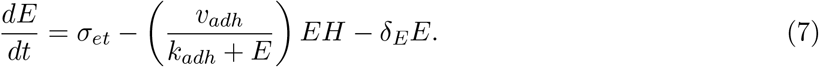

**Table 1:**
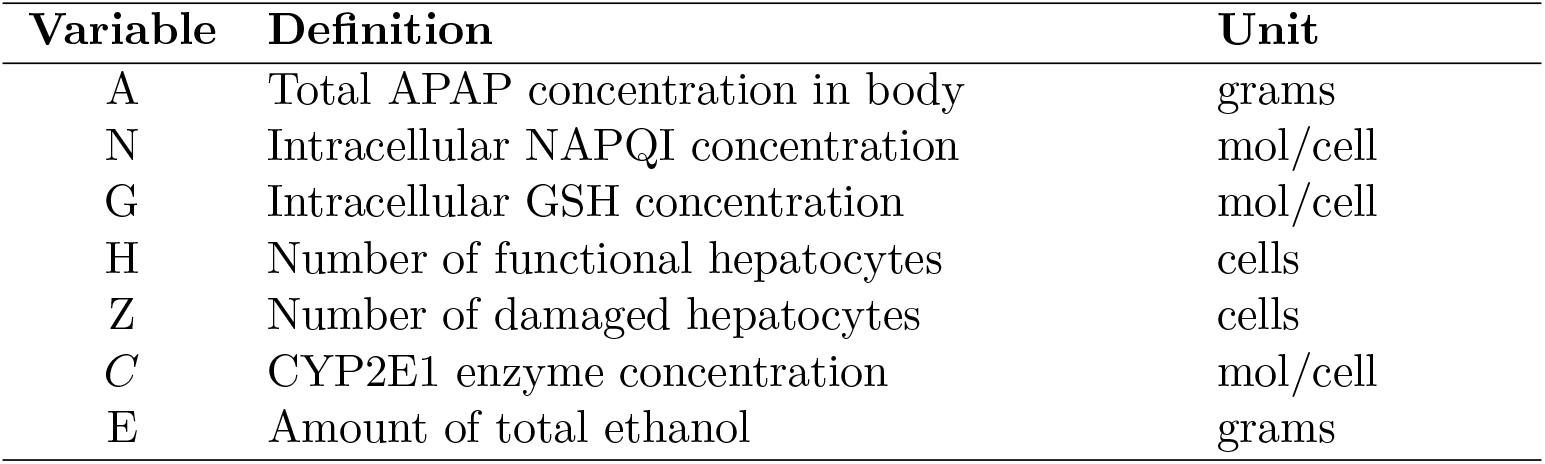
Variables used in the model

**Table 2:** Parameters used in the model

| Parameter | Description | Value and Reference |
| --- | --- | --- |
| $H_{max}$ | Number of hepatocytes in a healthy liver | $1.6 \times 10^{11}$ (Price and Alberti 1979, Sohlenius-Sternbeck 2006) |
| $\delta_Z$ | Damaged hepatocyte lysis rate | 5/day<br>Remien, Sussman, and Adler 2014 |
| $r$ | Functional hepatocyte regeneration rate | 1/day<br>(Furchtgott, Chow, and Periwal 2009) |
| $\eta$ | Functional hepatocyte damage rate | $5.12 \times 10^{13}$ cell/mol/day<br>Remien, Sussman, and Adler 2014 |
| $\alpha$ | APAP clearance rate by hepatocytes | 6.3/day<br>(Rumack and Matthew 1975) |
| $\delta_A$ | Unconjugated APAP clearance rate | .33/day<br>(J. Mitchell et al. 1973), (Bond 2009) |
| $p$ | Fraction of APAP oxidised to NAPQI | 0.05/day<br>(J. R. Mitchell et al. 1974, Bond 2009) |
| $q$ | Conversion factor from grams of APAP to moles of NAPQI | .0067 mol/gm<br>Remien, Sussman, and Adler 2014 |
| $\gamma$ | Functional hepatocyte damage rate | $1.6 \times 10^{18}$ cell/mol/day<br>(Miners and Kissinger 1979) |
| $\delta_G$ | GSH decay rate | 2/day<br>(Ookhtens et al. 1985),<br>(Lauterburg, Adams, and J. R. Mitchell 1984),<br>Aw et al. 1986) |
| $\kappa$ | GSH production rate | $1.375 \times 10^{-14}$ mol/cell/day<br>(Remien, Adler, et al. 2012) |
| $\delta_N$ | NAPQI clearance rate | $10^{-4}$/day<br>(Remien, Adler, et al. 2012) |
| $v_u$ | Maximum rate for APAP sulfation | 6gm/day<br>(Reith et al. 2009) |
| $k_u$ | Michaelis Menten constant for APAP sulfation | .0733gm (Reith et al. 2009) |
| $k_g$ | Glucoronidation rate of APAP | 6/day (Reddyhoff et al. 2015) |
| $k_{cyp}$ | Maximum rate of competitive inhibition by ethanol | .373/mole/day |
| $k_i$ | CYP2E1 enzyme dissociation constant | 200gm |
| $k_{deg}$ | Degradation rate of CYP2E1 enzyme | 2.273792/day (Chien, Thummel, and Slattery 1997) |
| $v_{adh}$ | Maximum rate for alcohol dehydrogenation | 210.16gm/day (Umulis et al. 2005) |
| $k_{adh}$ | Michaelis Menten constant for alcohol dehydrogenation | .04606gm (Umulis et al. 2005) |
| $\delta_E$ | Ethanol clearance rate | .035/day (Jones 2010) |
| $C_{01}$ | Normalized level of enzyme | 1 mol/cell (Chien, Thummel, and Slattery 1997) |
| $\sigma_{et}$ | Constant rate of alcohol ingestion | gm/day |

Before we study the effect of alcohol that causes change in APAP metabolism, we compare the existing model MALD of APAP metabolism in (Remien, Sussman, and Adler 2014) and its effect on liver injury with the current extended model where the effect of alcohol ingestion is taken into account. For, different overdose amount of APAP, we compare the NAPQI formation, the hepatocyte regeneration, glutathione production and APAP metabolism when alcohol is not ingested with APAP in the two models. Figures (2a), (2b),(2c) and (2d) show that the extended new model that includes the effect of alcohol ingestion gives us similar results as in the existing original model of MALD (see Remien, Sussman, and Adler 2014) when alcohol ingestion with single time APAP overdose is not considered.

**Figure 1:**
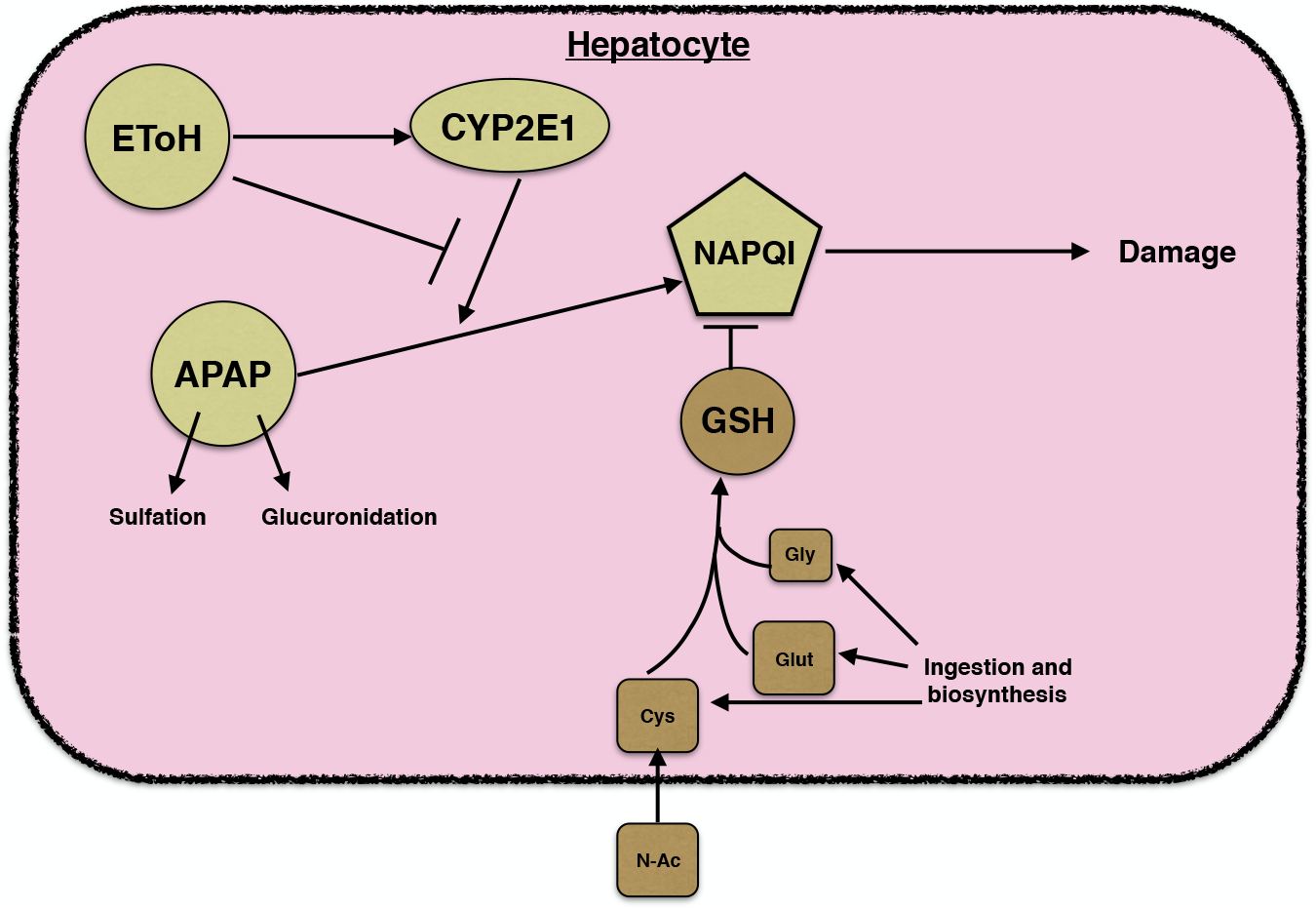
Schematic Diagram for the model dynamics within a hepatocyte.

**Figure 2:**
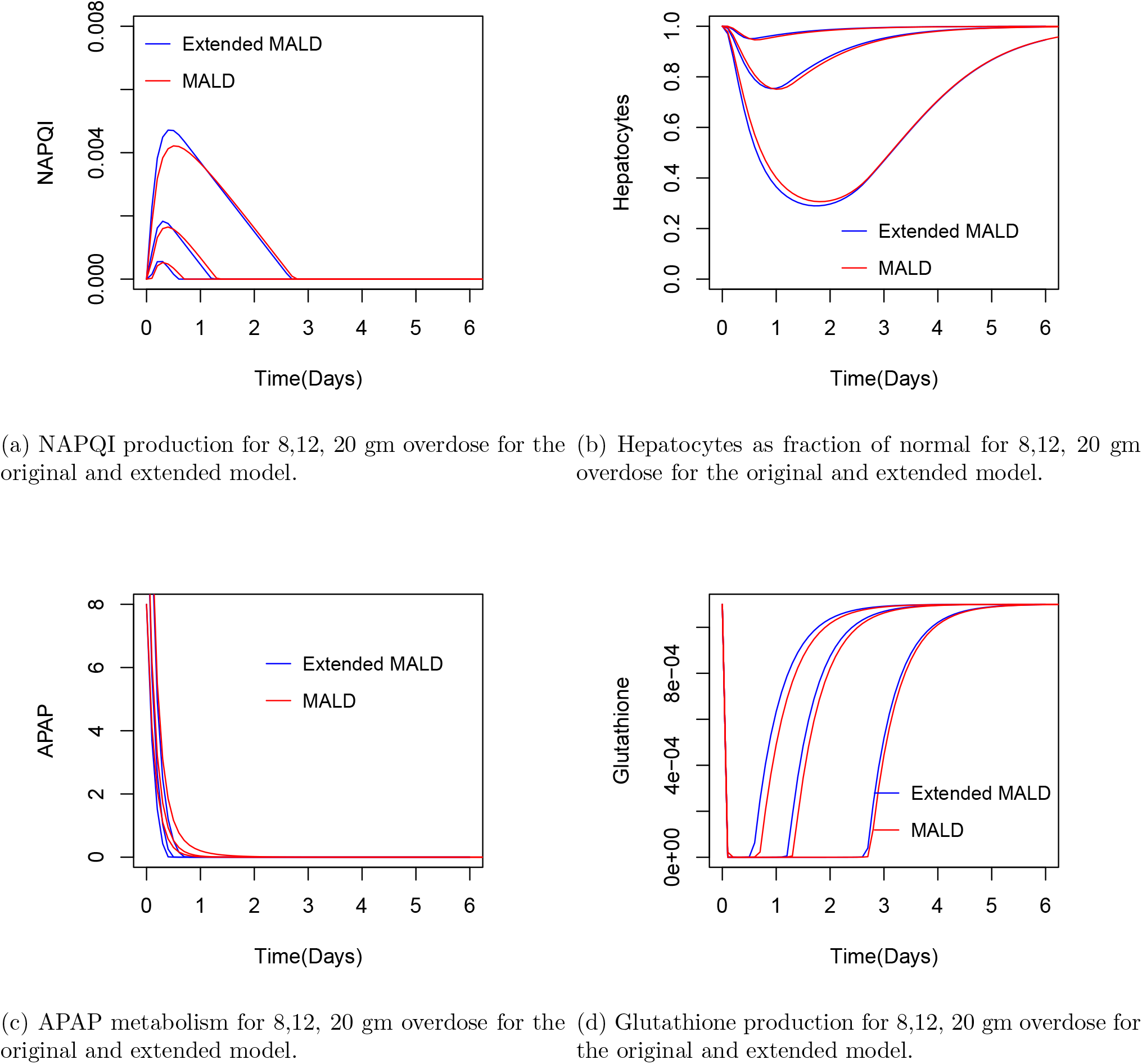
Time course for NAPQI, Hepatocytes as fraction of normal, APAP and GSH for the existing and the updated model without alcohol ingestion.

### 2.2 Steady States

We study the steady states in this section here to better understand chronic APAP metabolism in presence of alcohol in the body. We analyze the steady state of the model that includes chronic APAP metabolism and the effect of alcohol ingestion by setting the time derivative of the differential equation to zero. The chronic APAP metabolism along with alcohol ingestion model includes an APAP intake *σ* gm in the APAP equation 1. The steady state *A*\* of A, written in terms of the steady states of E, H and *C* is given by the positive root of the quadratic equation

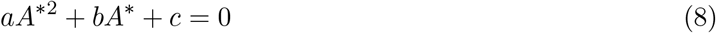

Where

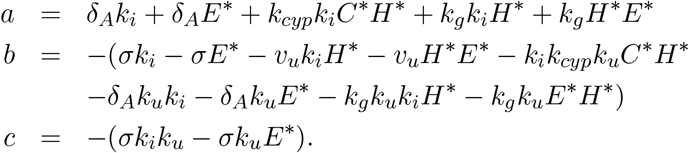

The steady state *G*\**, Z*\**, C*\* of G, Z and *C* are given in terms of the steady states *H*\*, *N* * of H and N respectively are

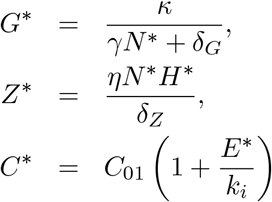

*E*\* the steady state of E is given by the positive root of the quadratic equation

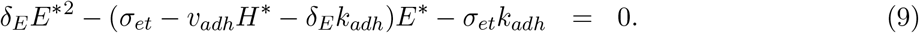

The steady state *H*\* of H in terms of the steady state *N* * of N are *H*\* = 0 and

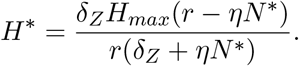

The case when *H*\* = 0, the steady states of A, E and *C* are given by

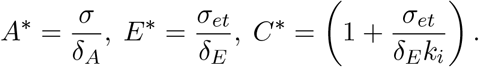

Hence the steady state of N corresponding to *H*\* = 0 is the positive root of the quadratic equation

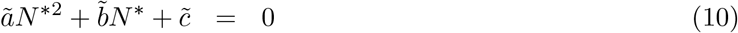

where where

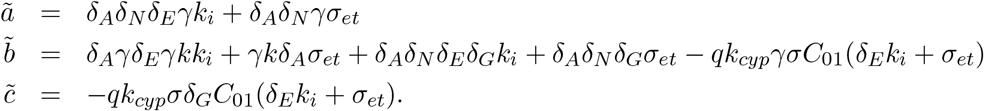

Equilibrium points corresponding to *H*\* = 0 and *H*\* = *H_max_* are given in Table 3. The steady state of N corresponding to 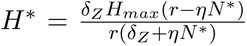 cannot explicitly be given as it involves the roots of *E*\* and *A*\*.

**Table 3:** Equilibrium points corresponding to *H*\* = 0 and *H*\* = *H*_*max*_

| Equilibrium points |  | Equilibrium points |  |
| --- | --- | --- | --- |
| $H^*$ | 0 | $H^*$ | $H_{max}$ |
| $Z^*$ | 0 | $N^*$ | 0 |
| $A^*$ | $\frac{\sigma}{\delta_A}$ | $Z^*$ | 0 |
| $E^*$ | $\frac{\sigma_{et}}{\delta_E}$ | $A^*$ | 0 |
| $C^*$ | $1 + \frac{\sigma_{et}}{\delta_E k_i}$ | $E^*$ | roots of (9) |
| $N^*$ | roots of (10) | $C^*$ | $1 + \frac{\sigma_{et}}{\delta_E k_i}$ |
| $G^*$ | $\frac{\kappa}{\gamma N^* + \delta_G}$ | $G^*$ | $\frac{\kappa}{\delta_G}$ |

#### 2.2.1 Stability of Steady States

We determine the stability of the steady states by analyzing the Jacobian of the system of differential equation. Linearizing the system of intracellular and hepatocyte differential equations yields the Jacobian

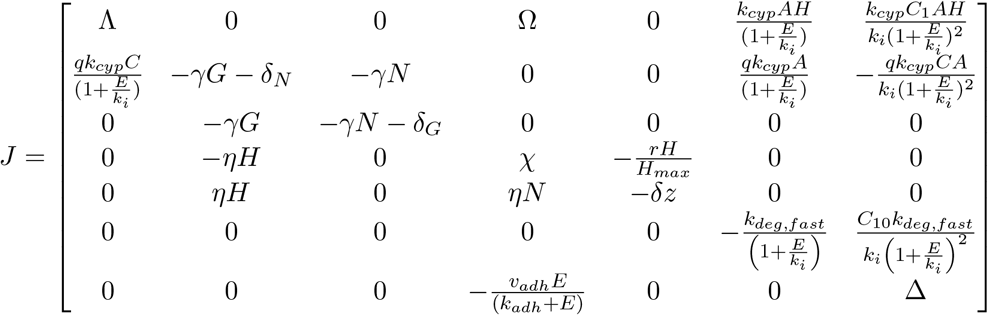

where 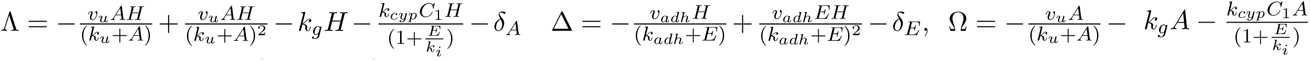*, and 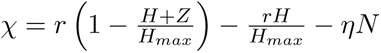.*

We analyse the steady states and the stability for the system corresponding to *H*\* = 0 and *H*\* = *H*_*max*_ numerically for the model that includes an APAP intake rate *σ* g/day and alcohol consumption *σ_et_* g/day in the body. The eigen values for the system corresponding to *H*\* = 0 and *H*\* = *H*_*max*_ are negative. We study the bifurcation analysis of this updated model with respect to the APAP intake rate *σ* g/day. We use the software XPPAUT AUTO to study the bifurcation analysis. We compare the bifurcation analysis with the original model MALD in (Remien, Sussman, and Adler 2014) in Fig(3a), Fig(3b), Fig(3c) and Fig(3d). *H*\* = 0 becomes stable at *σ*_1_ ≈ 0.3*g/day* through a transcritical bifurcation in both the cases. Corresponding to *H*\* ≈ *H*_*max*_, in the extended model, the stable fixed point and the unstable fixed point collides at *σ*_2_ *≈* 10.86*g/day* and vanish through a saddle node bifurcation while in the original model the point of collision of the stable and the unstable fixed point is *σ*_2_ *≈* 6.8*g/day*. Steady states at *H*\* ≈ *H*_*max*_ and *H*\* = 0 correspond to approximately no damage and full damage, respectively. The ratio of GSH and NAPQI production determines the point *σ*_2_. If the production of GSH is more than NAPQI then *σ* < *σ*_2_ and *σ* > *σ*_2_ when the production of NAPQI is more than GSH. The second case corresponds to a stable fixed point associated with full damage, for details (see ibid.).

**Figure 3:**
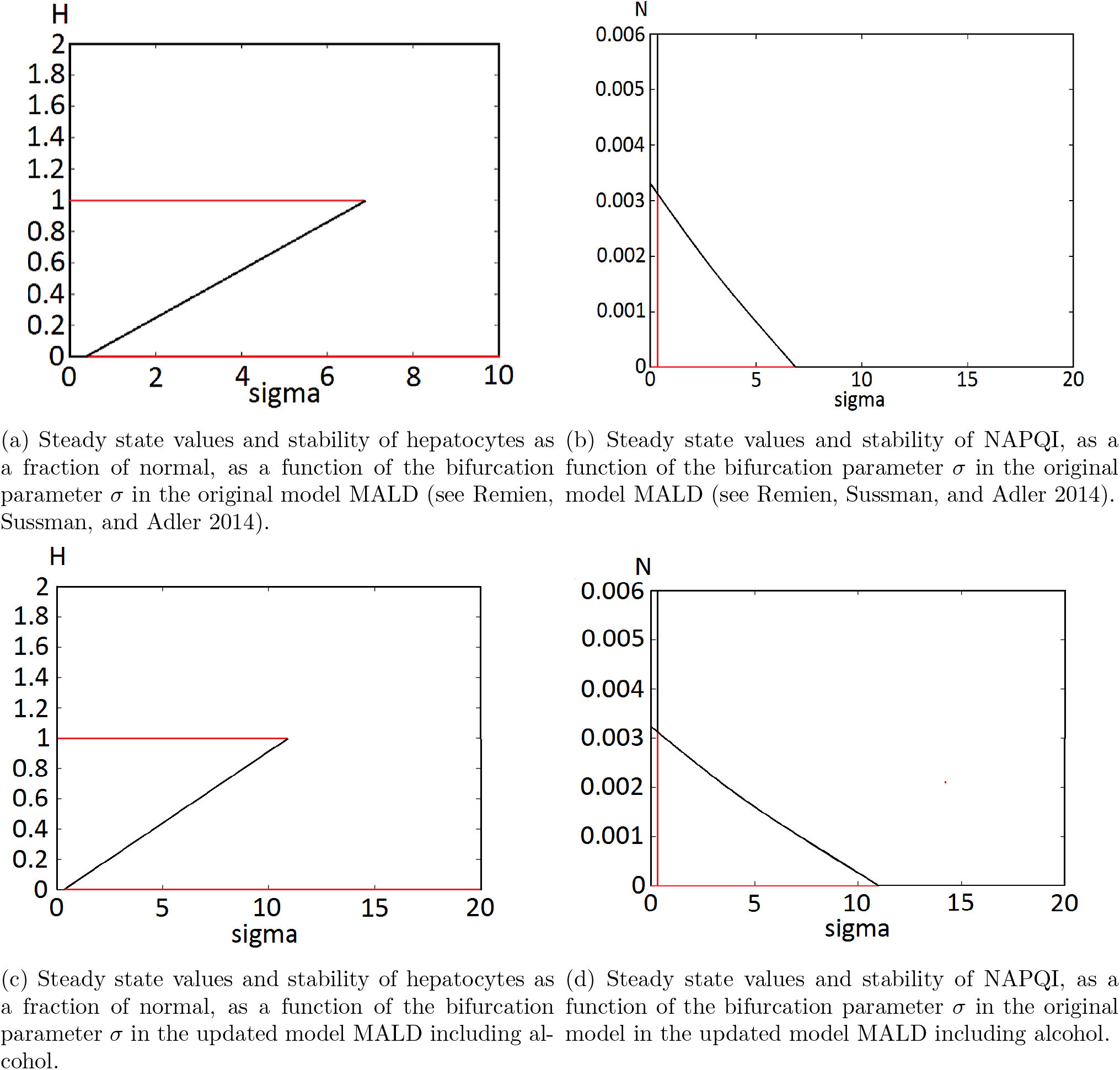
Bifurcation analysis of the original model MALD and the updated model with respect to the parameter *σ*.

## 3 Result

### 3.1 Acute Alcohol Ingestion

We study here how the model extended MALD including the effects of alcohol ingestion can help us estimate APAP induced liver injury based on the scenarios of acute use of alcohol. We consider here different time variant scenarios of alcohol and APAP ingestion. In the case of short term use of alcohol or acute use, we consider the cases

1. when alcohol is ingested within a range of one-seven days before APAP is consumed,
2. when APAP is simultaneously ingested along with alcohol
3. when alcohol is ingested within a range of 1-7 days after APAP is ingested.

If *t*_0_ is the time for single time APAP overdose and *t*\* is the time for alcohol intake then *t*\* = *t*_0_ for case 1, *t*\* = *t*_0_ for case 2 and *t*\* > *t*_0_ for case 3. We numerically solve our model given by eq(1) - eq(7) for a range of intake level of alcohol and size of overdose amount using R. The initial value for the CYP enzyme (C) is normalized to **1** and the initial value of NAPQI (N) and damaged hepatocyte (Z) are set at 0, healthy hepatocyte (H) set to maximum, Glutathione (G) set to steady state value.

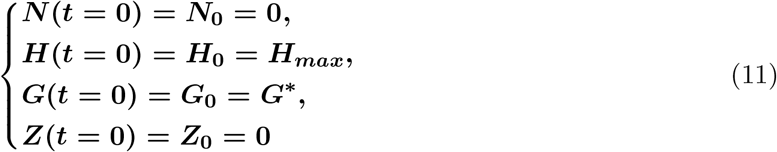

In the first case, we first numerically solve the subsystem eq(6) and eq(7) for *t* = *t*\* separately under the assumption *H* = *H*_*max*_ with initial condition *C*_0_ = 1 and for a range of one to seven days to obtain the value of the CYP enzyme levels of *C* for different amount of alcohol intake (*E*_0_). We then use these values of *C* as initial conditions for *C* and numerically simulate eq(1) - eq(7) to estimate the effect of liver injury caused when different level of alcohol is ingested for a range of one to seven days before APAP is taken. The initial condition for *C* is set to these previously obtained values and the initial condition for other variables are given by eq(11) to study the effect of this acute short term use of alcohol.

For the cases 2 and 3, when different doses of APAP ingestion simultaneously along with alcohol ingestion and when alcohol is ingested within a range of one to seven days after APAP is ingested, the initial condition for *C* is set to the no damaged state and the initial conditions for other variables are given by 11 to numerically track minimum hepatocyte level for different range of alcohol and APAP intake. We obtain the **minimum hepatocyte ratio** which is the ratio of the minimum hepatocyte count in the scenarios when alcohol is ingested with APAP to the minimum hepatocyte count when APAP is ingested only with no intake of alcohol at all.

Figure 4 shows that for different APAP overdose amounts and different alcohol amounts the minimum hepatocyte ratio which is the ratio of the minimum hepatocyte level in this updated model here to the minimum hepatocyte level in the original model MALD is less than or equal to one when alcohol is ingested **1** *−* **6** days before APAP is ingested causing more damage to the liver than the liver injury due to single APAP overdose. The minimum hepatocyte ratio is greater than or equal to one in the case of simultaneous ingestion and ingestion of alcohol a day prior to overdose APAP intake causing less damage or no damage to the liver than the one caused due to single APAP overdose. The minimum hepatocyte ratio for ingestion of alcohol after **1** – **7** days of APAP overdose is nearly one, showing that the consumption of alcohol a day after does not effect the liver injury already incurred. Shades of red indicate areas where there is more damage with alcohol ingestion than without, while shades of blue indicate area where there is less damage with alcohol. The model thus predicts short term ingestion of alcohol followed by a brief abstinence results in increased hepatoxicity when compared to an APAP overdose case for a nonalcoholic person. Thus the timing of alcohol ingestion is important in predicting liver injury in the case of APAP overdose.

**Figure 4:**
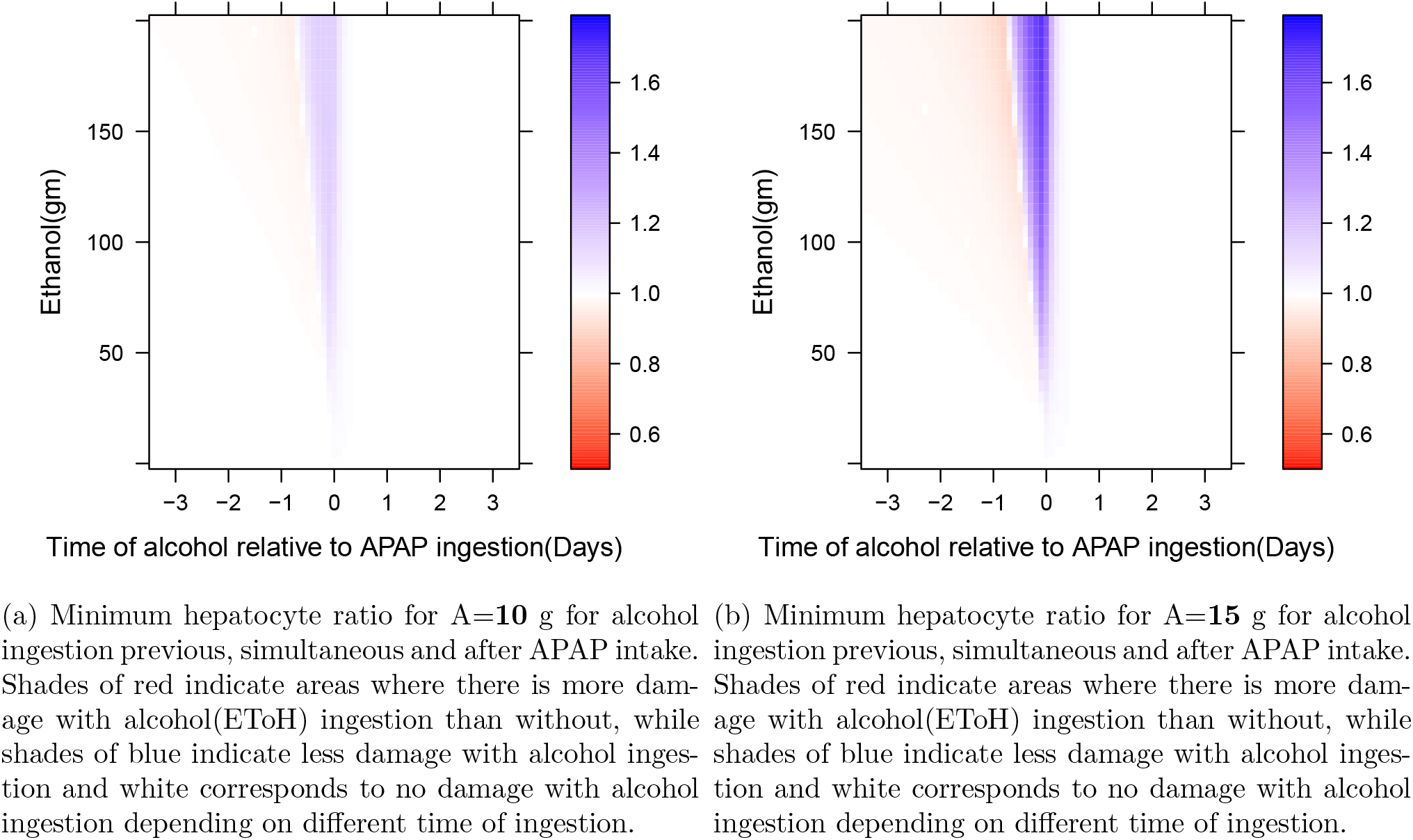
Minimum hepatocyte ratio which is the ratio of the minimum hepatocyte count with alcohol and APAP ingestion to the hepatocyte count for APAP ingestion only for the first three scenarios for a range of alcohol amount at different overdose amount of APAP intake.

### 3.2 Chronic Alcohol Ingestion

In the case of chronic use of alcohol, we consider the case when alcohol of different amount has been ingested for a considerable period of time and the enzymes *C* have reached a steady state due to periodic intake on a regular basis. Thus we consider the case when single overdose of APAP is ingested along with chronic use of alcohol. To study the extent of liver injury due to single APAP overdose against a background of chronic use of alcohol, we solve the steady state values of E numerically.

We then solve the steady state values of *C* from equation (6) for different amount of alcohol intake on a regular basis and use it as our initial condition for *C* when we estimate liver injury due to chronic alcohol ingestion. For different estimated overdose of APAP amount as *A*_0_ and different alcohol intake as *E*_0_ and with initial conditions in 11, we numerically solve the model eq(1) - eq(7) to obtain an estimated pre-treatment minimum hepatocyte level and outcome. Figures 5a, 5b show hepatocytes as fraction of normal for an overdose amount of **10**g and **15**g APAP for a range of different amount of chronic use of alcohol respectively. Figures 5c, 5d show the hepatocytes as fraction of normal for a range of overdose amount for a chronic use of alcohol of **50***g* and **100**g respectively. The chronic use of alcohol on a single time APAP overdose shows us an increase in hepatoxicity in the liver as the amount of alcohol intake gets higher. Thus our synthetic data verifies the hypothesis that alcohol consumption increases susceptibility to APAP toxicity in the scenario of chronic intake of alcohol (Bray et al. 1991). Susceptibility to toxicity increases with higher amount of alcohol intake thus increasing the risk of liver injury.

**Figure 5:**
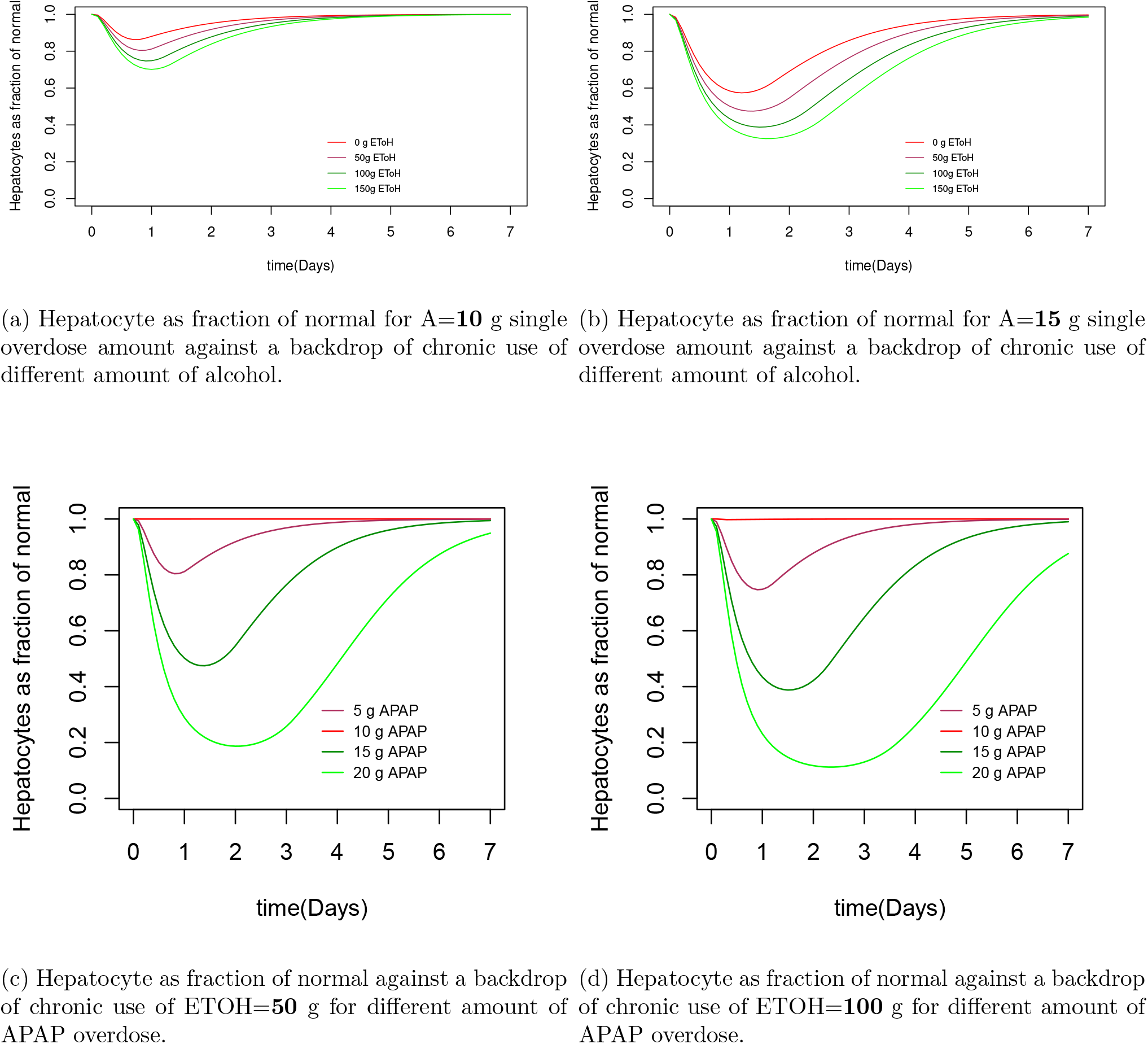
Hepatocytes as fraction of normal level for different amount of APAP overdose and for a range of different amount of chronic alcohol intake.

## 4 Discussion

We extend the mathematical model of acetaminophen induced liver (MALD) injury to include the dual effect of alcohol induction and inhibition. We explicitly model the effect of alcohol on metabolism of APAP by CYP2E1 enzymes and induction of CYP21 enzymes by alcohol. The timing of alcohol ingestion relative to APAP is crucial in determining liver damage. The results here show that the time of ingestion of alcohol with respect to APAP even in the backdrop of acute ingestion of alcohol have different effect on the liver. The risk of APAP induced hepatotoxicity is increased if APAP is ingested shortly after alcohol is cleared from the body. This increased susceptibility to APAP hepatotoxicity is successfully reflected in our mathematical model. The time course of the hepatocyte injury for different amount of alcohol intake gives us an estimate of the liver injury. Our model also shows the protective effect of acute consumption of alcohol when APAP is ingested in presence of alcohol in body. This is due to the fact that alcohol acts both as a substrate and as an inhibitor of CYP2E1 (see Thummel et al. 2000; Slattery, Nelson, and Thummel 1996). Our results on the liver injury of single time APAP overdose along with alcohol ingestion at different time scenarios relative to the overdose matched with the observation of clinical studies that shows that simultaneous ingestion of alcohol and APAP and intake of alcohol after APAP overdose for non-alcoholic individuals results in lesser production of NAPQI with decreased risk of hepatotoxicity (see Chien, Thummel, and Slattery 1997). Single time APAP overdose in the scenario of acute and chronic ingestion of alcohol is also numerically simulated here.

The time course of liver injury in the chronic scenario of alcohol ingestion for a range of alcohol intake is easily understood from our model and is in sync with clinical reports. While (Chien, Thummel, and Slattery 1997; Lesser, Vietti, and Clark 1986; Banda and Quart 1982) has modeled alcohol metabolism and induction of CYP enzymes by alcohol, here we have added the effect of alcohol on APAP metabolism by CYP enzyme to determine mathematically how it effects the metabolism and measure the resulting liver injury. The model gives an insight of the extent of liver injury for range of alcohol ingestion amount and timing relative to APAP by the synthetic data simulated through the mathematical model here. This model in future can help us also determine the risk factors associated with APAP induced liver injury for alcoholic patients by fitting this model with patients data containing alcohol information along with other bio-markers. This model can be tested in a multicenter dataset in future to validate the results of the model.

